# Instant preparative droplet reaction assisted by shaking

**DOI:** 10.1101/2025.04.30.651440

**Authors:** Ming Bi, Lang Zhao, Zhixin Tian

## Abstract

Instant micro- and macro-preparative droplet reaction assisted by hand shaking (shake reaction) was found and observed, where reactions were carried out inside aqueous droplets dispersed in an organic solvent. This shake reaction was successfully benchmarked with both organic reactions of protein reduction by dithiothreitol and alkylation by iodoacetamide as well as biological reaction of protein digestion by enzyme trypsin. The reaction time of shake reaction was substantially shortened from the traditional bulk solution reactions, specifically 1 min for reduction (from 20 min), 1 min for alkylation (from 30 min), 5 min for digestion (from 960 min). The high efficiency of shake reaction comes from the micro-nano droplet formation assisted by shaking. Shake reaction can arguably be extended to any liquid-phase inorganic, organic and biological reactions from micro- to macro-preparative scale, and thus undoubtedly find wide applications in both academic research and industry production.

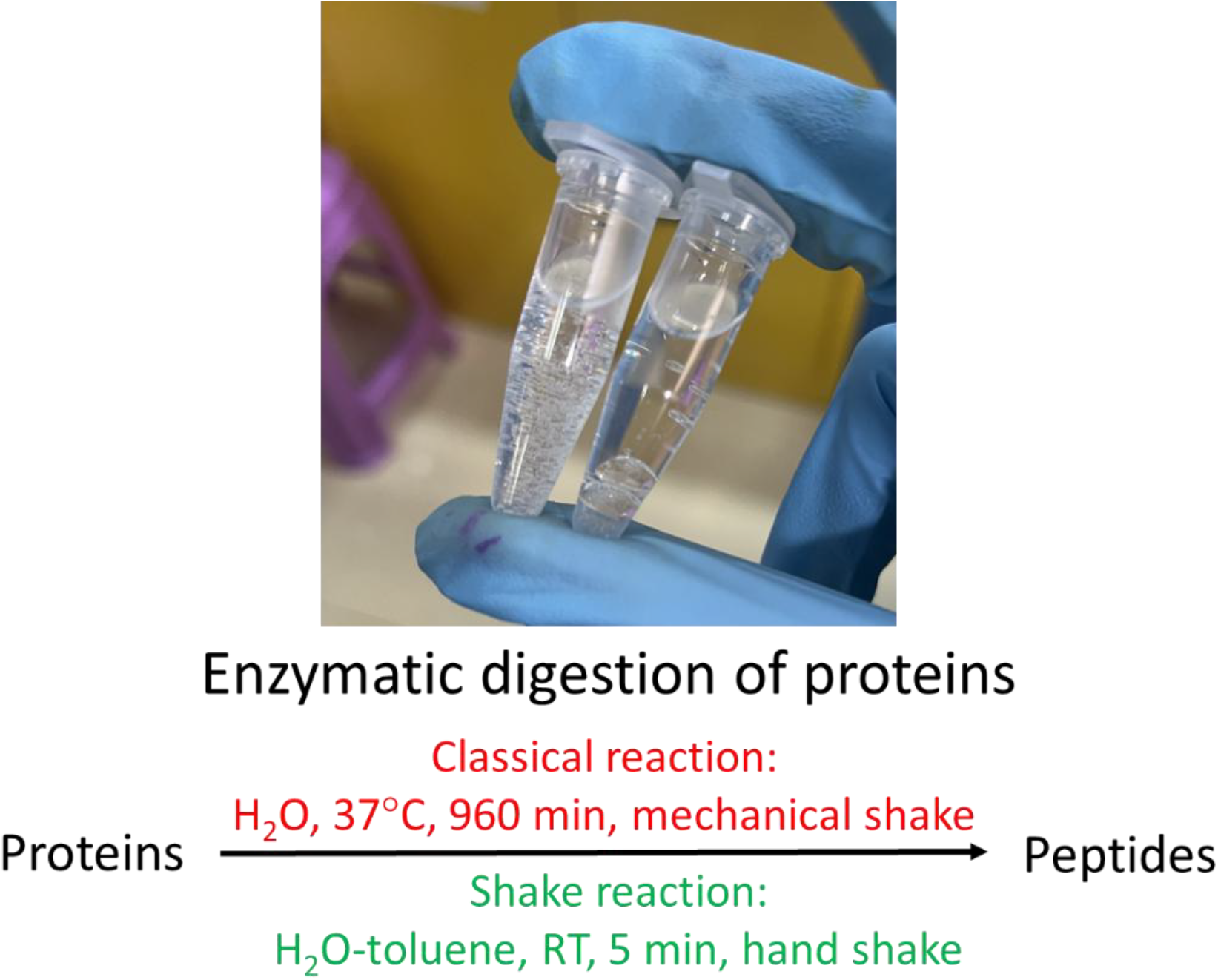

## INTRODUCTION

Droplets have been reported to be efficient reaction vessels (reactors) in the cases of isolated droplets inside the microfluidic channel(*1*), on-the-fly droplets among the sonic spray ionization stream(*2*), acoustically levitated droplets generated from field-induced droplet ionization(*3, 4*), as well as droplets on the artificial Teflon-coated hydrophobic surface in the study of droplet-based one-pot preparation for proteomic samples (DROPPS)(*5*). In the exemplary reaction of protein digestion by trypsin, micro-droplet reaction on-the-fly in sonic spray ionization had similar digestion specificity of NIST 8671 mAb antibody in terms of semi- and unspecific-cleaved peptides with the overnight (14 hrs) bulk digestion at 37°C, the milli-droplet DROPPS reaction time at 37°C with mixing at 500 rpm decreased the reaction time to 2 hrs from overnight (16 hrs) of the bulk reaction; the former (with smaller droplet size) is faster but only suitable for the analytical purpose, the latter (with larger droplet size) is slower but suitable for the micro-preparative purpose. Thus, fast micro- (even nano-) droplet reaction covering the micro- to macro-preparative scale remains to be further investigated.

Here we report instant micro- and macro-preparative droplet reaction assisted by hand shaking (shake reaction), where reaction takes place in aqueous droplets dispersed in an organic solvent. This shake reaction was successfully benchmarked with both organic reactions of protein reduction by dithiothreitol and alkylation by iodoacetamide as well as biological reaction of protein digestion by enzyme trypsin. The reaction time of shake reaction was substantially shortened from the traditional bulk solution reactions, specifically 1 min for reduction (from 20 min), 1 min for alkylation (from 30 min), 5 min for digestion (from 960 min). The high efficiency of shake reaction comes from the micro-nano droplet formation assisted by shaking. Reactions with potential hazards or gas release can be carried out with alternative automated mechanical (instead of hand) shaking. Shake reaction can arguably be extended to any liquid-phase inorganic, organic and biological reactions from micro- to macro-preparative scale, and thus undoubtedly find wide applications in both academic research and industry production.

## EXPERIMENTAL

### Materials and reagents

All reagents including dithiothreitol (DTT), iodoacetamide (IAA), acetonitrile (ACN) and formic acid (FA) were obtained from Sigma-Aldrich (St. Louis, USA). Sequencing grade modified trypsin was purchased from Promega (V5111, Madison, USA). SDS, Tris, urea, ammonium bicarbonate (ABC) and BCA Protein Assay Kit were purchased from Sagon Biotech (Shanghai, China). Ultrapure water was produced onsite with a Millipore Simplicity System (Billerica, MA, USA). The tissue protein sample of this study is the same with our previous study(*6*), and is used in all the QC, shake reaction and bulk reaction.

### Organic reactions of protein reduction by dithiothreitol and alkylation by iodoacetamide

The liver tissue sample were stored in –80°C until used. After homogenized in SDS/TRIS (HCl) solution at a ratio of 1:20 (v/v) along with a cocktail and shaking for 30 min, the resulting mixture was then centrifugated at 14,000 rpm for 30 min, and the supernatant was collected and added to ice-cold acetone for precipitation in −20 °C for 3 h. The protein pellets, from centrifugation at 10,000 g for 10 min were then redissolved in 8 M urea and diluted with 50 mM ABC until the urea concentration was reduced to below 1 M. resulting in a concentration of approximately 1.5 μg/μL. For the shake reaction, 70 μL protein solution was added into 1 mL toluene dispersant followed by the addition of 7 μL of 100 mM DTT to the protein solution and was shook 1 min and centrifuge 20 s to aggregate dispersed droplets together. Based on this approach, the alkylation reaction was completed by added 7 μL of 200 mM IAA for 1 min and 7 μL of 100 mM DTT for 1 min to consuming redundant IAA.

### Biological reaction of protein digestion by enzyme trypsin

For the shake digestion, 0.1 μg/μL trypsin solution were added at different enzyme/protein ratio for various reaction time. Digestion reactions were quenched by 10 μL of 10 % FA solution. For the normal digestion, the protein solution (about 500 μg) was mixed with trypsin (1:50, w/w) in approximately 1 mL 50 mM ABC solution and incubated 16 h with gentle shaking at 37 °C.

The digested peptides were then desalted using C18 (5 mg, 15 μm, Jupiter, Phenomenex) solid phase extraction (SPE) columns. After equilibrated with 80 % ACN containing 0.1 % TFA and subsequently 0.1 % TFA, the desalting SPE column were loaded in digestion solution for 5 times for sufficient binding. After 8 times wash by 0.1 % TFA, peptides were eluted by 50 % ACN containing 0.1 % TFA and 80 % ACN containing 5 % TFA. The eluants were combined, dried in a Speed-Vac (Thermo Fisher Scientific, USA).

### RPLC-MS/MS analysis of the digested peptides

RPLC-MS/MS analysis of the digested peptides was carried out on a Dionex Ultimate 3000 RSLC nano-HPLC system coupled online through nano-ESI with a Q-Exactive mass spectrometer (Thermo Fisher Scientific, San Jose, CA, USA). Buffer A is 0.1% FA, aqueous solution, and Buffer B is 0.1% FA, 90 % ACN solution. Chromatographic columns packed with C18 (5 μm, Jupiter, Phenomenex) were composed of trap column (5 cm long, 200 μm i.d) and analytical column (60 cm long, 75 μm i.d). The multi-step gradient settings were as follows: 12 % buffer B 10 min, 20–45 % buffer B 95 min, 45–98 % buffer B 5 min, 98 % buffer B 3 min, 98-2 % buffer B 2 min, 8 % buffer B 5 min.

MS spectra were acquired in the 350-1800 *m/z* range using a mass resolution 70 k (*m/z* 200). For MS/MS spectra, the mass resolution was set at 17.5 k. Fragmentation was obtained in a data-dependent mode (Top30) with higher-energy collisional dissociation (HCD). The automatic gain control (AGC) target value and maximum injection time were placed at 1×10^6^ and 50 ms for MS and at 5×10^5^ and 250 ms for MS/MS scans. Isolation window and dynamic exclusion were set at 2 *m/z* and 30.0 s. Stepped normalized collision energies was optimally set at 30.0%. The temperature of the ion transfer capillary was set to 250 °C. The spray voltage was set to 2.8 kV. QC of the sensitivity of the LC-MS/MS analysis was monitored throughout the experiments by analysis of pre-prepared peptide sample.

### Bioinformatic identification of peptides

Database search of the raw LC-MS/MS datasets and identification of peptides and proteins was carried out in FragPipe (version 22.0). The adopted mass tolerance for the precursor and fragment ions are 10 and 20 ppm; acetylation at the protein N-terminal, oxidation on methionine as well as carbamidomethylation on cysteine (for assessment of the reaction efficiencies of reduction and alkylation) are set as dynamic modifications. False discovery rate (FDR) was controlled to be no more than 1%.

Each LC-MS analysis was carried out with three technical replicates (if not described otherwise), and average number of peptide and protein IDs with relative standard deviation were reported. Statistical distribution of peptide length, percentage of cysteine alkylation, miss cleavages and confidence score of hyperscore was carried out for assessment of identification confidence and reaction efficiency.

## RESULTS

### QC of the sensitivity of the LC-MS/MS analysis

A complex tissue peptide sample was adopted for the monitoring and maintenance of the sensitivity of the LC-MS/MS analysis as tracked by the number of peptide and protein IDs with average and RSD of 6500±2.5% and 1117±4.3%, respectively (**Supplemental Figure S1**).

### Shake reaction in the biological reaction of protein digestion by enzyme trypsin

With the reaction temperature of room temperature (RT, 25°C), protein concentration of 1.5 μg/μL, dependence of reaction rates as characterized by total number of peptide and protein IDs (the same thereafter) of the biological reaction of protein digestion by enzyme trypsin on the amount of enzyme, reaction time, and volume of the organic phase were investigated.

For the dependence of the protein digestion on the amount of enzyme, with the temperature of RT, the aqueous volume of 91 μL (the summed volume of the solutions of protein, DTT, IAA) and 1 mL toluene and reaction time of 5 min, the trypsin/protein ratio (w/w) of 1:20, 1:50 and 1:100 was investigated (**Figure 1**). With the trypsin/protein ratio goes from 1:100 to 1:20, 1:50 is optimal, although the numbers of peptide IDs of 1:20 is slightly higher (6850±894 vs. 6667±304) and the ratio of zero miss cleavage is slightly higher (58.3% vs. 51.5%). The distribution of cysteine alkylation The ratio of 1:50 is chosen for the following experiments.

**Figure 1.**
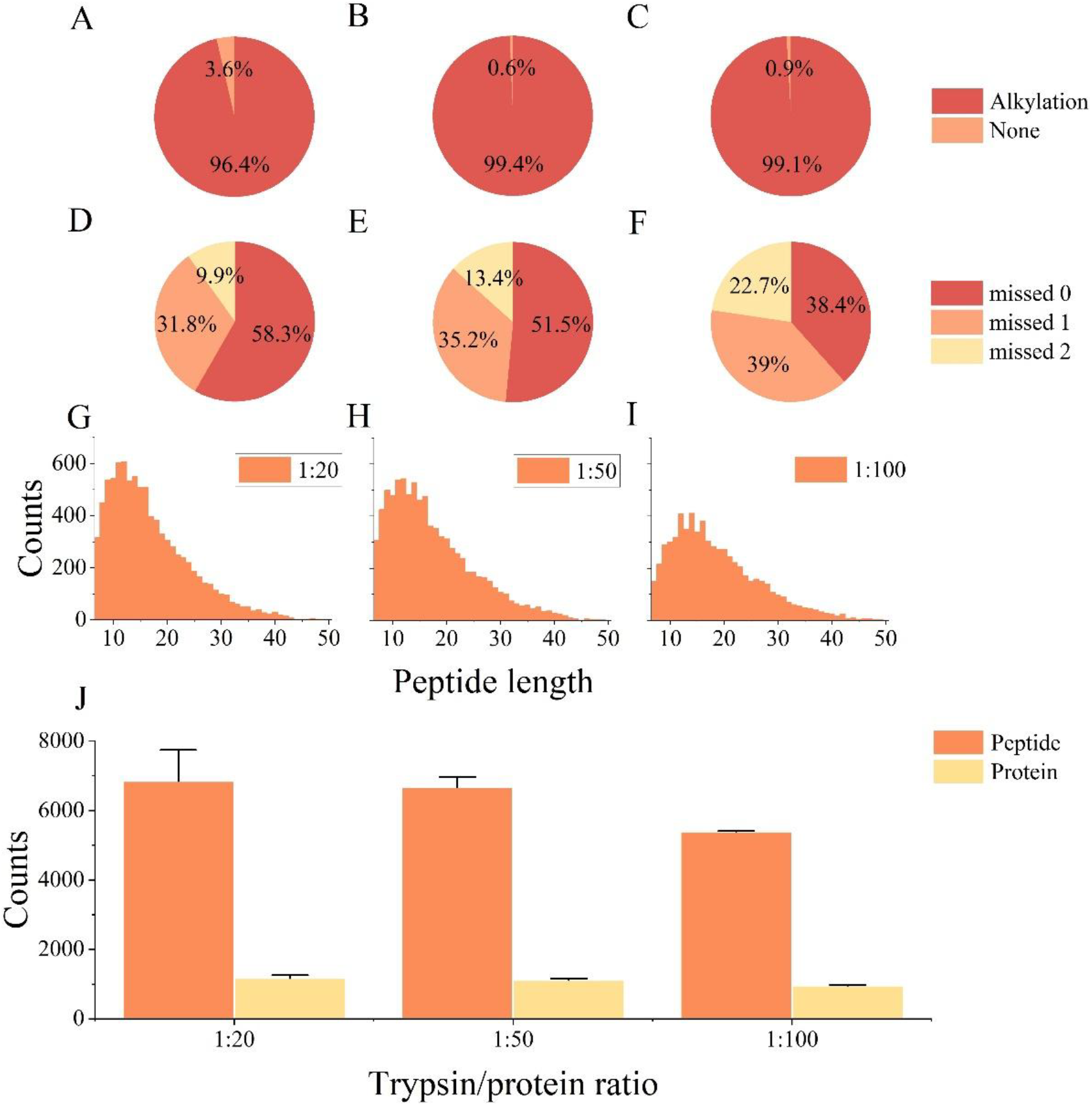
Influence of the amount of enzyme of 1:20 (A, D, G, J), 1:50 (B, E, H, J) and 1:100 (C, F, I, J) in the shake reaction of biological enzymatic protein digestion by trypsin with the reaction time of 5 min; room temperature, 1 mL toluene and hand shaking; (A, B, C) cysteine alkylation, (D, E, F) miss cleavages (G, H, I) peptide length distribution, (J) peptide and protein IDs.

For the dependence of the protein digestion on the reaction time, with the protein/enzyme ration 50:1, RT, toluene volume of 1 mL, the reaction time of 2.5 min, 5 min and 10 min was investigated (**Supplemental Figure S2**). 5 min is optimal in terms of number of IDs and the percentage of zero miss cleavage, and is chosen for the following experiment.

For investigation of the possibility of replacing hand shaking with automated mechanical shaking in the case of reactions with potential hazards or not suitable for hand shaking (such as with gas release), with the trypsin/protein ratio of 1:50 (w/w), the reaction time of 5 min, room temperature and 1 mL toluene volume, protein digestion with automated vortex shaking was carried out (**Supplemental Figure S3**). Similar performance of hand vs. machine shaking in terms of the number of peptide IDs (6667±304 vs. 6186±41) and the percentage of zero miss cleavage (51.5% vs. 41.7%) was observed.

For the dependence of protein digestion on the organic **dispersant**, with the trypsin/protein ratio of 1:50, the reaction time of 5 min, room temperature, 1 mL dispersant and hand shaking, toluene and fluoro oil were comparatively investigated (**Supplemental Figure S4**). Toluene is much better than the fluoro oil; alternative hexane, heptane and isooctane were also investigated, but none is better than toluene (data not shown).

For the exploration of the shake reaction for macro-preparative production, the reaction volume was scaled from 1 mL to 10 mL (while keeping the concentrations of all the reactants the same, **Supplemental Figure S5**), the IDs of peptides are 6667±304 vs. 5329±211 and the percentage of zero miss cleavage is 51.5% and 37.3%, respectively, with somewhat decrease of the reaction efficiency. The reason maybe due to the influence of un-digested proteins in the de-salting process, which guarantees further improvement space.

For comparison of the reaction efficiency of shake reaction with the bulk reaction at the optimal conditions of each reaction (**Figure 2**), similar number of peptide IDs (6667±304) and protein IDs (6625±109) were obtained; however, the percentage of cysteine alkylation of the shake reaction (99.4%) is slightly higher than that of the bulk reaction (91.7%), while the percentage of zero miss cleavage of the shake reaction (51.5%) is slightly lower than that of the bulk reaction (60.3%). The overlap of peptides and proteins are ca. 70% and 80%; for the shared peptides, the confidence score of peptide identification (hyperscore) of the shake reaction is better than the bulk reaction. The distribution of the peptide length and hyperscore is similar.

**Figure 2.**
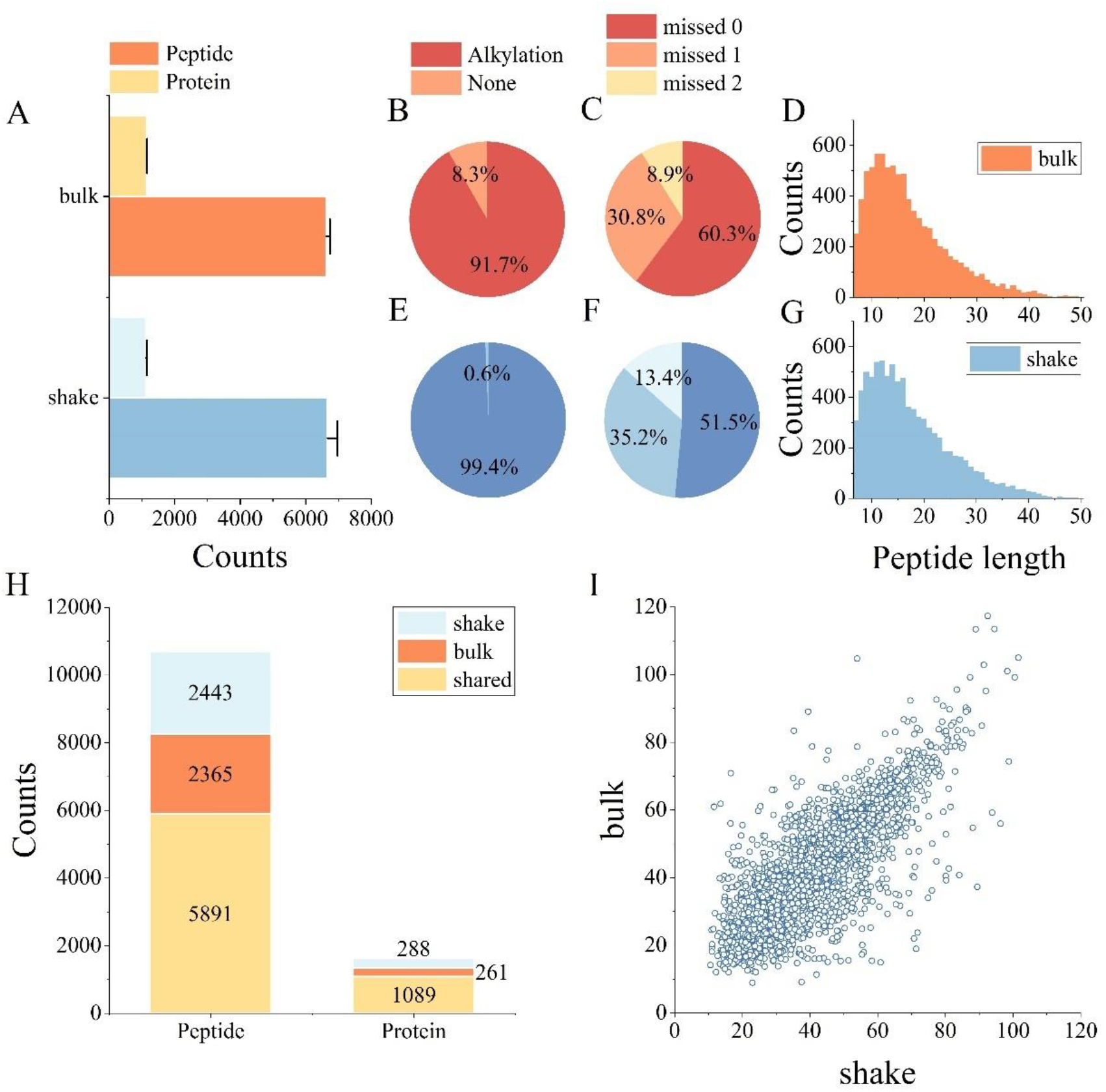
Efficiency comparison of the shake reaction (A, E, F, G, H, I) with the bulk reaction (A, B, C, D, H, I) in the biological enzymatic protein digestion by trypsin; (A) peptide and protein IDs, (B, E) cysteine alkylation, (C, F) miss cleavages, (D, G) peptide length distribution, (H) overlap of the peptide and protein IDs, (I) hyperscore of the shared peptide IDs.

It should be noted that in addition to the shake reaction reported here, pressurization has been a common way of accelerating protein enzymatic digestion, and one of the latest report(*7*) is “at 45,000 psi for 60 cycles (20 s at 45,000 psi and 10 s at ambient pressure per cycle)”, i. e, 30 min of reaction time in total, with the specialized equipment of PCT MicroPestles (PBI Pressure BioSciences, Inc.).

### Shake reaction in the organic reactions of protein reduction and alkylation

Parallel and proportional to the decrease of reaction time of protein digestion, the reaction time of protein reduction by DTT and alkylation by IAA in shake reaction was decreased to 1 min (from 20 min of regular bulk reaction) and 1 min (from 30 min of regular bulk reaction), respectively. These two reactions were complete as demonstrated by the percentages of cysteine alkylation (99.4%) which was set at a dyanmic modification during the database search (**Figure 1B**).

## DISCUSSION

For the absolute peptide and protein IDs, the numbers can be substantial bigger if newer mass spectrometers (such as Thermo Astral, Bruker timsTOF, Sciex x600 zenoTOF) are used.

For the formation of aqueous nano-macro droplets in the organic phase, emulsifier is not always necessary under continuous shaking (either hand or automated mechanical), but for the protein solution in this study, 50 mM ABC has been found to be necessary. A higher concentration of ABC (500 mM) leads to smaller droplets and better dispersion, but the identified peptides and proteins are smaller (**Supplemental Figure S6**), which may be due to inhibition of trypsin activity by the basic solution, and intermediate concentrations are worth further investigation. Sodium dodecyl sulfate (SDS), the common, surfactant, was found to be not a good emulsifier for the protein solution to form uniform size droplets in toluene. The match of a reactant and its suitable emulsifier for shake reaction needs further investigation and possibly to be customized (i.e., one reactant, one emulsifier).

## CONCLUSIONS AND PERSPECTIVES

Efficient shake reaction through dispersion of the reaction phase as nano-micro droplets in an immiscible liquid phase was discovered and successfully benchmarked with both organic reactions of protein reduction by dithiothreitol and alkylation by iodoacetamide as well as biological reaction of protein enzymatic digestion by trypsin. The reaction time was decreased by hundreds of fold, the reaction temperature was decreased to the convenient room temperature, and the need of pressurizing, heating and shaking equipment (at the micro-preparative scale in a centrifuge tube) is also diminished. Shake reaction is proposed to be capable of being extended to other organic, biological as well as inorganic reactions at the micro-preparative scale, and thus undoubtedly find wide applications in both academic research and industry production.

## AUTHOR CONTRIBUTIONS

ZXT conceived the study; MB did the experiments and data analysis, LZ provided and assisted with the experimental equipment and instruments, ZXT wrote the manuscript.

## ACKNOWLEDGMENTS

The authors are grateful to Prof. Keqi Tang and Dr. Hanzhong Hu at Ningbo University for providing the tims TOF instrument for some initial experiments. This research was financially supported by the National Natural Science Foundation of China (22074105) and the Shanghai Science and Technology Commission (14DZ2261100).

## DATA AVAILABILITY

The raw LC-MS/MS datasets are available at ProteomeXchange Consortium (https://www.proteomexchange.org/).

## SUPPLEMENTAL INFORMATION

**Supplemental Figure S1.**
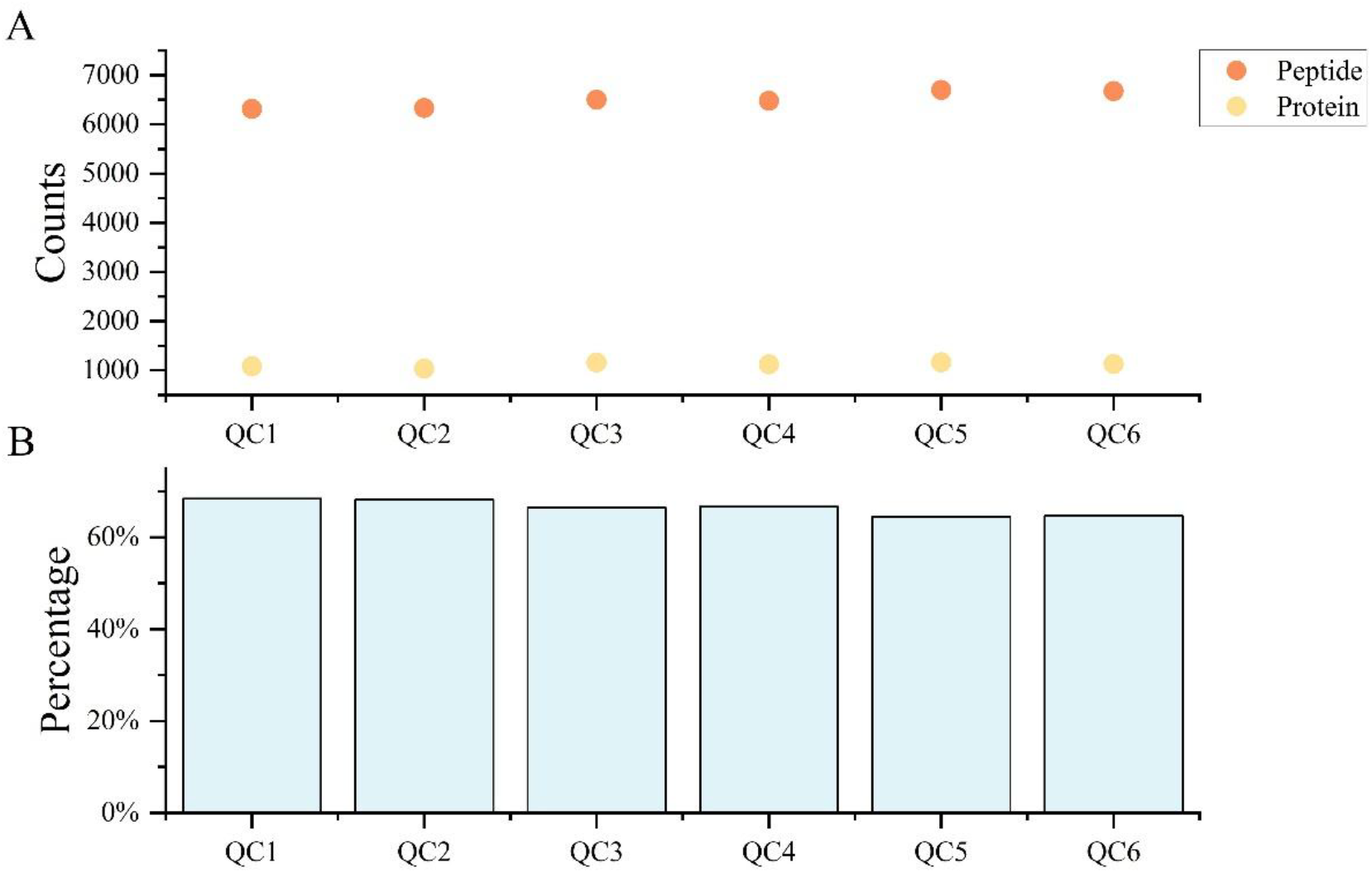
QC of the sensitivity of the LC-MS/MS analysis covering the whole period of the experiments in this study.

**Supplemental Figure S2.**
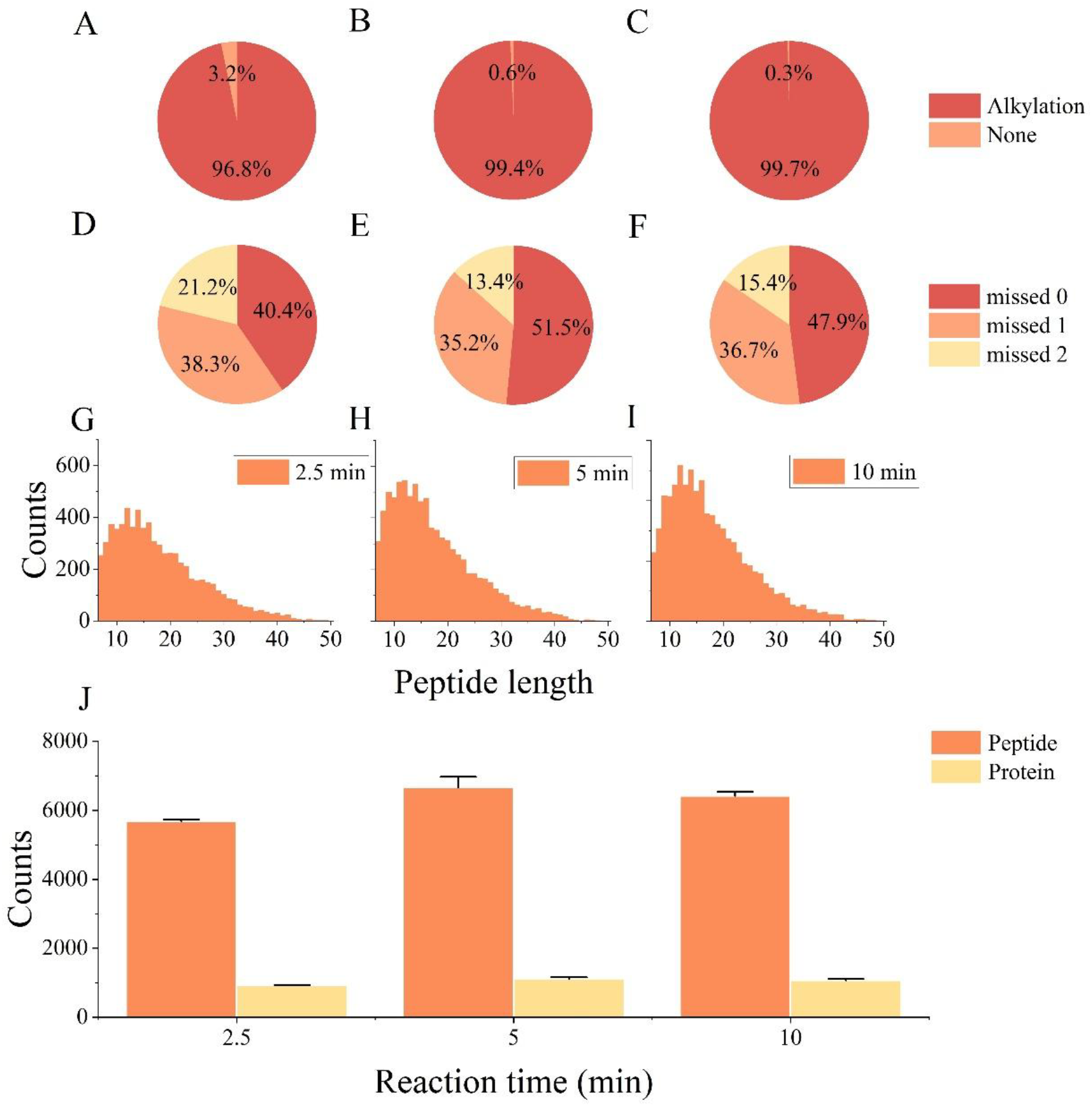
Influence of the reaction time of 2.5 min (A, D, G, J), 5 min (B, E, H, J) and 10 min (C, F, I, J) in the shake reaction of biological enzymatic protein digestion by trypsin with the trypsin/protein ratio of 1:50 (w/w), RT and 1 mL toluene; (A, B, C) cysteine alkylation, (D, E, F) miss cleavages (G, H, I) peptide length distribution, (J) peptide and protein IDs.

**Supplemental Figure S3.**
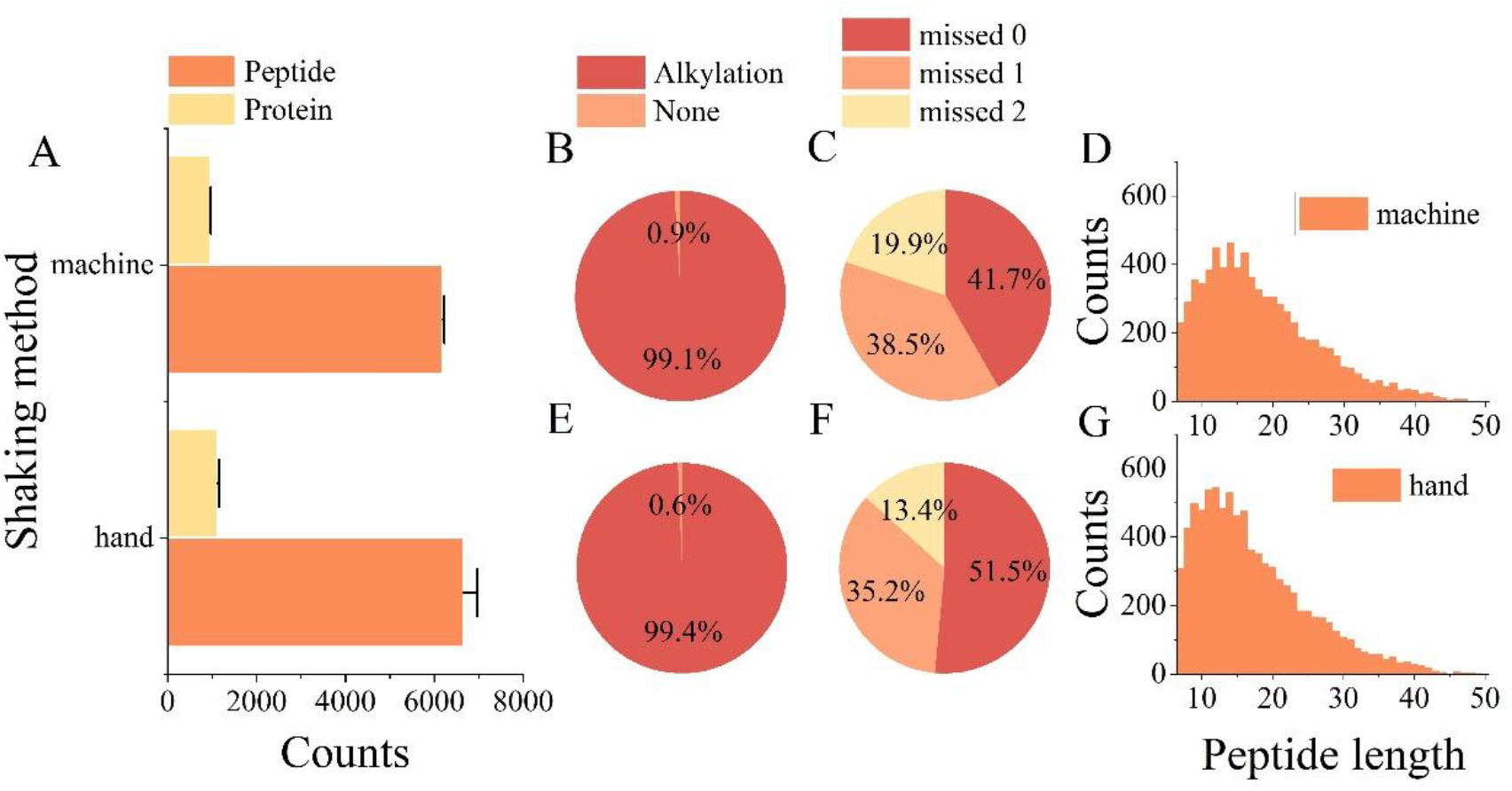
Influence of the shaking method of hand (A, E, F, G) vs. vortex (A, B, C, D) in the shake reaction of biological enzymatic protein digestion by trypsin with enzyme/protein ratio of 1:50 (w/w), reaction time of 5 min, RT and 1 mL toluene. (A) peptide and protein IDs, (B, E) cysteine alkylation, (C, F) miss cleavages, (D, G) peptide length distribution.

**Supplemental Figure S4.**
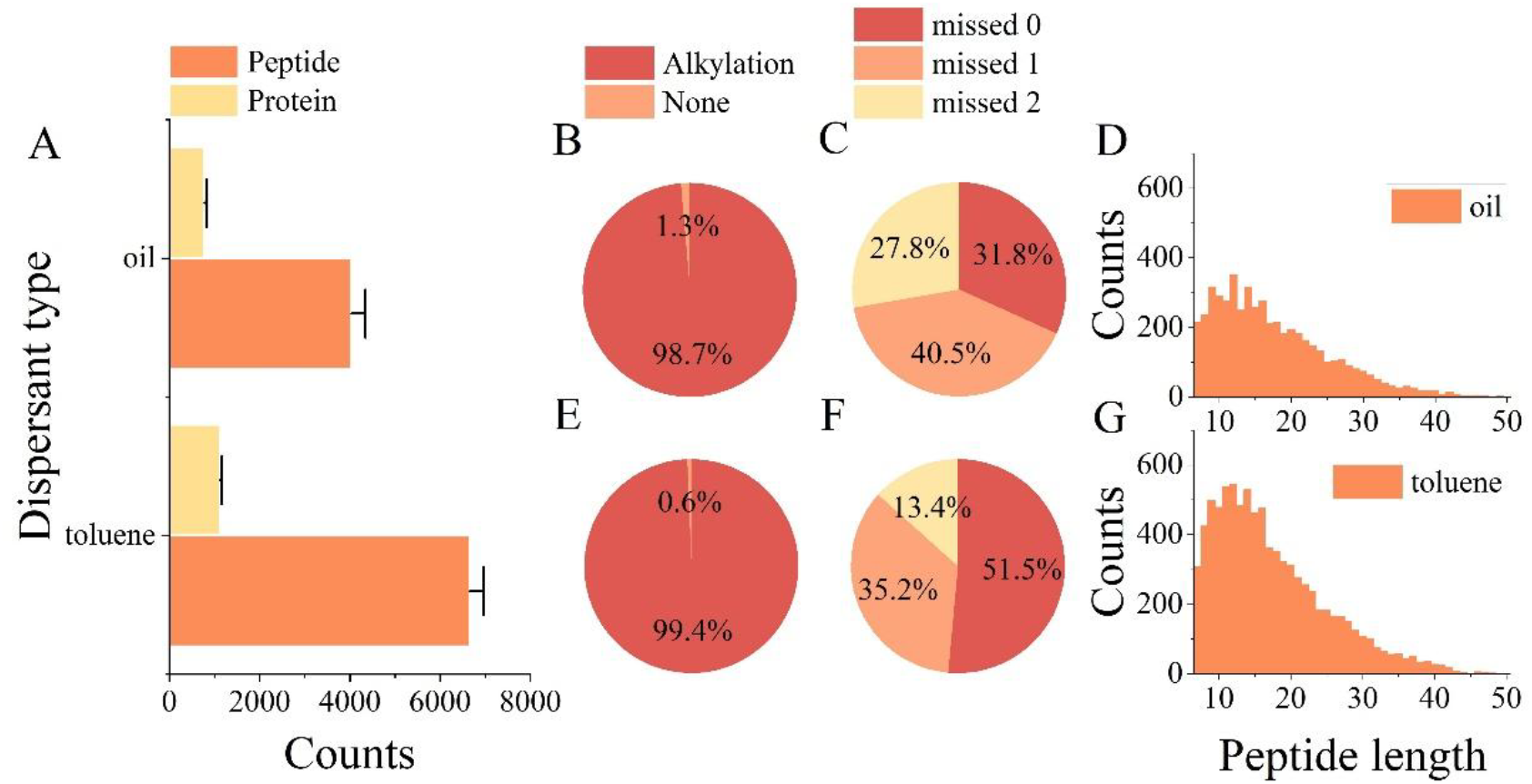
Influence of **dispersants** of toluene (A, E, F, G) vs. fluoro oil (A, B, C, D) in the shake reaction of biological enzymatic protein digestion by trypsin with the enzyme/protein ratio of 1:50, reaction time of 5 min, room temperature, 1 mL dispersant and hand shaking; (**A)** peptide and protein IDs, (**B, E**) cysteine alkylation, (**C, F**) miss cleavages, (**D, G**) peptide length distribution.

**Supplemental Figure S5.**
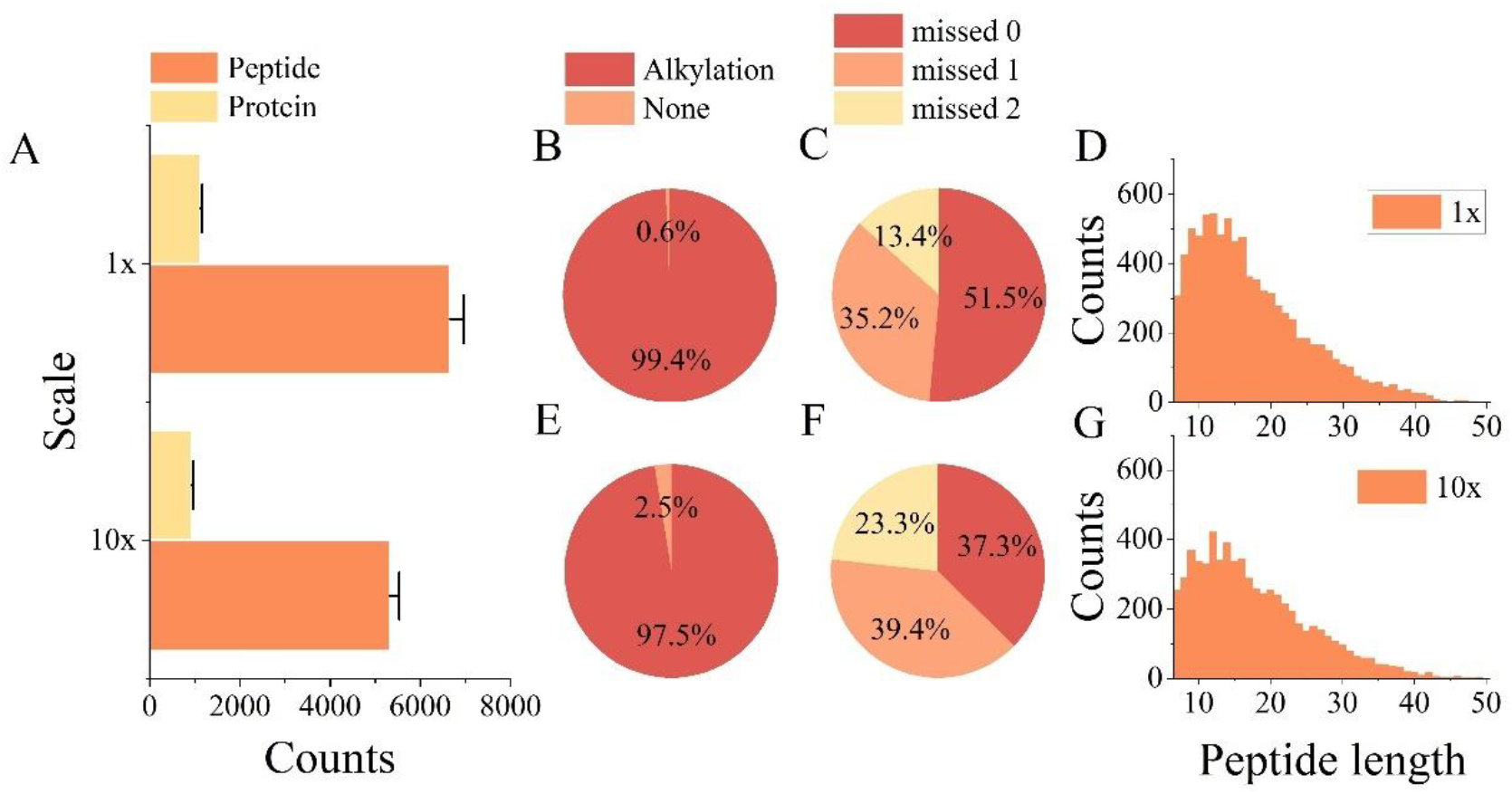
Scale-up of the reaction volume from 1 mL (A, B, C, D) to 10 mL (A, E, F, G) in the shake reaction of biological enzymatic protein digestion by trypsin with the enzyme/protein ratio of 1:50, reaction time of 5 min, room temperature and hand shaking; (**A)** peptide and protein IDs, (**B, E**) cysteine alkylation, (**C, F**) miss cleavages, (**D, G**) peptide length distribution.

**Supplemental Figure S6.**
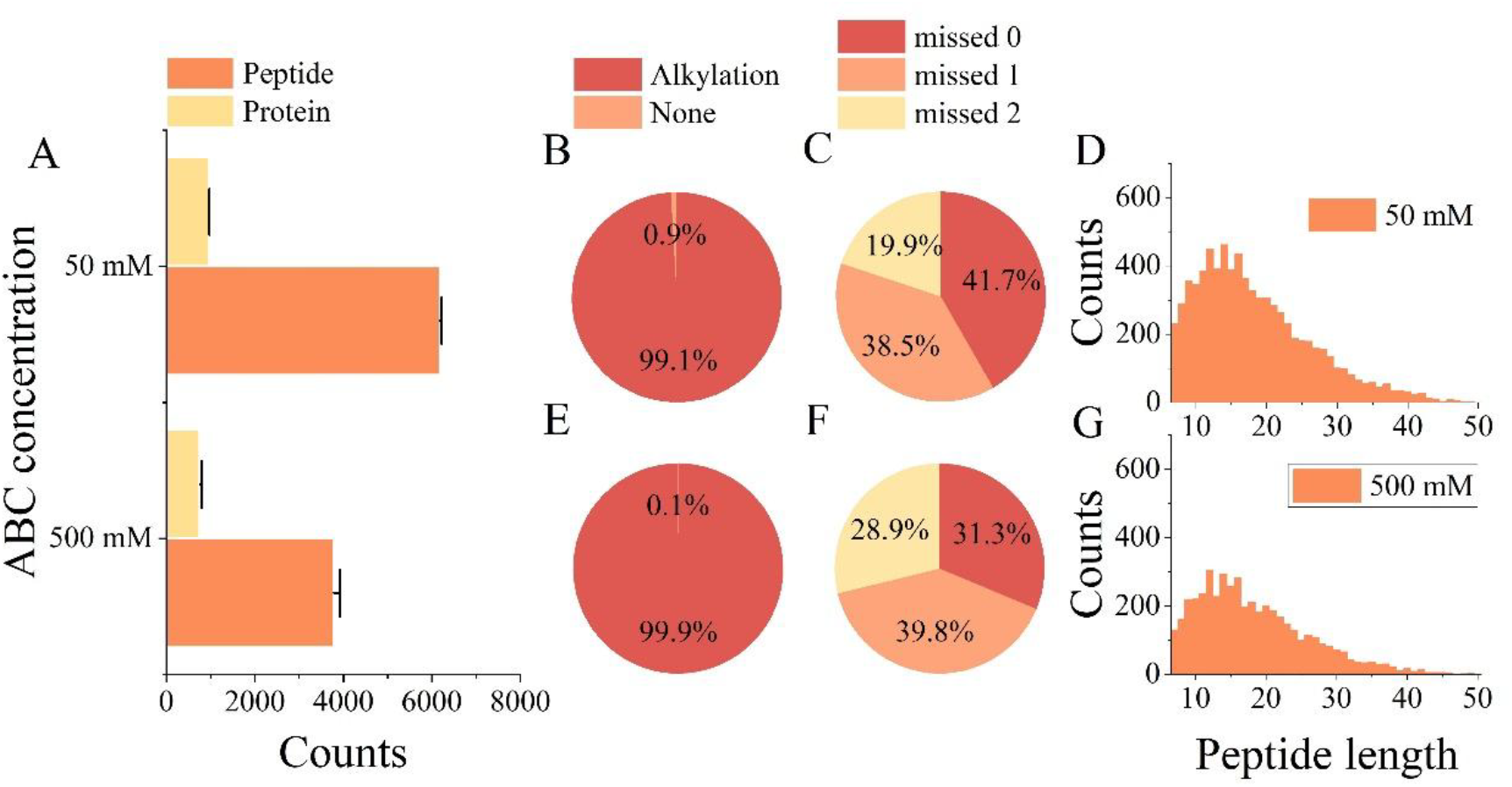
Influence of the concentration of ammonium bicarbonate of 50 mM (A, B, C, D) vs. 500 mM (A, E, F, G) in the shake reaction of biological enzymatic protein digestion by trypsin with the enzyme/protein ratio of 1:50, reaction time of 5 min, room temperature, 1 mL toluene, and vortex horizontal shaking; (**A)** peptide and protein IDs, (**B, E**) cysteine alkylation, (**C, F**) miss cleavages, (**D, G**) peptide length distribution.

